# Sensitivity of dairy calf *Salmonella enterica* serotype Cerro isolates to infection-relevant stressors

**DOI:** 10.1101/2024.01.10.575057

**Authors:** Sarah M. Raabis, Trina L. Westerman, Eddy Cruz, Courtney L. Deblois, Garret Suen, Johanna R. Elfenbein

## Abstract

*Salmonella enterica* serotype Cerro (*S.* Cerro) is an emerging *Salmonella* serotype isolated from cattle, but the association of *S.* Cerro with disease is not well understood. While comparative genomic analyses of bovine *S.* Cerro isolates have indicated mutations in elements associated with virulence, the correlation of *S.* Cerro fecal shedding with clinical disease in cattle varies between epidemiologic studies. The primary objective of this study was to characterize the infection-relevant phenotypes of *S.* Cerro fecal isolates obtained from neonatal calves born on a dairy farm in Wisconsin, USA. The *S.* Cerro isolates varied in biofilm production and sensitivity to the bile salt deoxycholate. All *S.* Cerro isolates were sensitive to sodium hypochlorite, hydrogen peroxide, and acidic shock. However, *S.* Cerro isolates were resistant to nitric oxide stress. Two *S*. Cerro isolates were unable to compete with *S.* Typhimurium during infection of calf ligated intestinal loops, indicating decreased fitness *in vivo*. Together, our data suggest that *S.* Cerro is sensitive to some innate antimicrobial defenses present in the gut, many of which are also used to control *Salmonella* in the environment. The observed phenotypic variation in *S.* Cerro isolates from a single farm suggest phenotypic plasticity that could impact infectious potential, transmission, and persistence on a farm.

**Importance:** *Salmonella enterica* is a zoonotic pathogen that threatens both human and animal health. *Salmonella enterica* serotype Cerro is being isolated from cattle at increasing frequency over the past two decades, however its association with clinical disease is unclear. The goal of this study was to characterize infection-relevant phenotypes of *S.* Cerro isolates obtained from dairy calves from a single farm. Our work shows that there can be variation among temporally-related *S.* Cerro isolates and that these isolates are sensitive to killing by toxic compounds of the innate immune system and those used for environmental control of *Salmonella*. This work contributes to our understanding of the pathogenic potential of the emerging pathogen *S*. Cerro.

## Introduction

Non-typhoidal *Salmonella enterica* are a global threat to human health, with more than 150 million cases and over 50,000 deaths reported annually (1). Alongside the risk to food safety, non-typhoidal *Salmonella enterica* are an important cause of disease and decreased productivity in both calves and adult dairy cattle (2, 3). During the last two decades, *Salmonella enterica* serotype Cerro has emerged as one of the most common serotypes isolated from both clinically ill and apparently healthy cattle in the United States (https://www.cdc.gov/nationalsurveillance/pdfs/salmonella-serotypes-isolated-animals-and-related-sources-508.pdf). In Wisconsin, the prevalence of *S*. Cerro isolations increased from less than 1% in 2006 to 37% in 2015 of all *Salmonella-*positive bovine samples submitted to the Wisconsin Veterinary Diagnostic Laboratory (4). The Minnesota Veterinary Diagnostic Laboratory also reported an increase in *S.* Cerro-positive isolates from 6.6% of all *Salmonella* isolates recovered from cattle in 2006 to over 20% in 2013 (5). In Pennsylvania, the prevalence of *S.* Cerro from bovine-associated *Salmonella*-positive samples increased from 14.3% in 2005 to 36.1% in 2010 (6). Thus, the rate of *S*. Cerro isolation is increasing in numerous geographical regions in the United States.

Despite its emergence in cattle, *S.* Cerro rarely causes disease in humans, and its association with clinical disease in cattle is not well characterized (7). Results from an epidemiological study of dairy herds in New York showed that cattle with clinical signs consistent with salmonellosis were 3.9 times more likely to be *S.* Cerro positive than apparently healthy cattle (2). Another study evaluating *Salmonella* shedding patterns in Ohio dairy herds found that *S*. Cerro was isolated from some adults with low milk production and intermittent enteritis, and from calves with severe diarrhea on one of the dairies (8). However, in one Pennsylvania dairy herd, 75% of the herd was shedding *S.* Cerro for at least six months but clinical salmonellosis was not observed in cattle shedding *S*. Cerro during the study period (9). Given that the majority of *S.* Cerro infections documented in cattle are subclinical, a hypothesis has emerged that this serotype has reduced virulence capacity (10).

Comparative analyses of bovine *S*. Cerro isolates have demonstrated mutations, deletions, and reduced expression of genes associated with virulence when compared with *S. enterica* serotypes that commonly cause food-borne disease in humans. Most *S*. Cerro isolates sequenced to date have a premature stop codon in *sopA* (10–12). SopA is a key type-3 secretion system-1 (T3SS1) effector protein that works along with SopB, SopD, SopE2 and SipA to cause diarrhea during *Salmonella* Typhimurium infection (13, 14). In addition to carriage of a mutation in *sopA*, some *S*. Cerro isolates have reduced expression of T3SS1 proteins when compared with *S*. Typhimurium or *S*. Dublin (15, 16). Consistent with the genomic and transcriptomic observations of alterations to the T3SS1, *S*. Cerro isolates are less invasive in tissue culture than other serotypes (17). Other notable loci which are absent or contain gene deletions in some *S*. Cerro isolates include *Salmonella* Pathogenicity Island (SPI)-10 (absent), deletions in SPI-6 and -13 (homologs of *STM0293* and *STM3123*), deletion or nonsense mutations in a thiosulfate reductase gene (*phsA*) in the tetrathionate reduction pathway, and deletion of a D-alanine transporter important for intracellular survival (homologs of *STM1633-STM1637*) (10, 11). Many of these loci in *S*. Typhimurium support virulence and colonization of the mammalian host (18–21). Although genomics studies infer reduced virulence of *S.* Cerro for mammals, there is a lack of data exploring the phenotypes of *S.* Cerro isolates in infection-relevant conditions.

The central objective of this study was to characterize the infection-relevant phenotypes of four *S.* Cerro isolates obtained from one-day-old dairy calves born within a period of 30 days. Our work demonstrates phenotypic variation within *S*. Cerro isolates and enhanced sensitivity to antibacterial killing mechanisms that are employed by the innate immune system and used for environmental control of *Salmonella*.

## Results

### Variability of disease severity in S. Cerro-colonized neonatal calves

Calves were removed from the dairy farm within 2 hours of birth and were housed in individual pens isolated from all other animals to eliminate exposure to infectious diseases. Clinical observational data and results of screening fecal samples for *Salmonella* spp. are listed in Table 1. Passive transfer of immunity was fair to excellent in all four calves (22). All calves were colonized by *S.* Cerro at 1 day of life and continued to shed the organism until at least 10 days of life. The meconium was negative for *Salmonella* spp. for the two calves in which it was tested. During the 11-day monitoring period, all calves had at least 1 day of pyrexia (rectal temperature ≥ 102.5°F). Calves 5 and 6 had multiple days of diarrhea, defined as a fecal score >1, while calves 3 and 4 had no evidence of diarrhea (Table 1). No animal required medical intervention for either pyrexia or diarrhea. Calves 3 and 4 had *S.* Cerro isolated from cecal tissue, ileal contents and mesenteric lymph nodes at postmortem examination. Colon contents of calf 5 was positive for *Salmonella* spp. but no *Salmonella* spp. was isolated from calf 6 at postmortem examination. These observational data suggest that calves were exposed to *S*. Cerro during or immediately after parturition and demonstrate that *S*. Cerro can disseminate to and persist within mesenteric lymph nodes of neonatal calves for at least 11 days.

**Table 1.** Clinical findings and results of *Salmonella* testing from naturally infected neonatal calves.

| Clinical variable | Calf #3 | Calf #4 | Calf #5 | Calf #6 |
| --- | --- | --- | --- | --- |
| sex | bull | heifer | bull | heifer |
| date of birth | 4/14/21 | 4/15/21 | 5/4/21 | 5/7/21 |
| STP (g/dL) | 5.7 | 6.2 | 5.1 | 5.2 |
| Number of days febrile ($T \geq 102.5$) | 1 | 1 | 3 | 1 |
| Number of days with diarrhea (FS $\geq 2$) | 0 | 0 | 3 | 4 |
| **Salmonella fecal screening** |  |  |  |  |
| Day 0 | N/D | N/D | negative | negative |
| Day 1 | S. Cerro | S. Cerro | S. Cerro | S. Cerro |
| Day 2 | N/D | positive | positive | positive |
| Day 3 | N/D | N/D | positive | positive |
| Day 4 | positive | positive | positive | positive |
| Day 6 | N/D | S. Cerro | positive | negative |
| Day 7 | S. Cerro | N/D | N/D | N/D |
| Day 10 | positive (*invA* +) | positive (*invA* +) | positive (*invA* +) | positive (*invA* +) |
| **Postmortem Examination** |  |  |  |  |
| Postmortem age | Day 12 | Day 11 | Day 16 | Day 13 |
| gross lesions | MLN lymphadenopathy | MLN lymphadenopathy | none observed | colonic mucosal edema |
| **Tissue culture** |  |  |  |  |
| MLN | S. Cerro | S. Cerro | negative | negative |
| Ileum contents | positive (*invA* +) | negative | negative | negative |
| Ileum tissue | negative | negative | negative | negative |
| Cecum contents | negative | negative | negative | negative |
| Cecum tissue | positive (*invA* +) | positive (*invA* +) | negative | negative |
| Colon contents | negative | negative | positive (*invA* +) | negative |
| Colon tissue | negative | negative | negative | negative |
| liver | negative | negative | negative | negative |
| gall bladder | N/D | N/D | negative | negative |
| spleen | negative | negative | negative | negative |
| right subiliac LN | negative | negative | negative | negative |
| left subiliac LN | negative | negative | negative | negative |

\*N/D – not tested; positive – *Salmonella* spp. identified by culture; positive (*invA*+) – *Salmonella* spp. identified by culture and isolate verified by PCR; S. Cerro – isolate serotyped; negative – no *Salmonella* spp. isolated; STP – serum total protein; FS – fecal score; MLN – mesenteric lymph node; LN – lymph node

### *S.* Cerro *in vitro* growth, motility, and biofilm formation

We first sought to establish the *in vitro* growth characteristics of *S*. Cerro isolated from feces of one-day old calves. S. Cerro isolates were named for the calf from which they originated (e.g. strain C3 was isolated from calf 3). The *S.* Cerro strains were isolated from calves born within one month of each other on the same farm, so we hypothesized that growth kinetics would be similar in various *in vitro* conditions. We tested aerobic growth in rich and minimal media and anaerobic growth in rich media to mimic the full variety of oxygen tensions along the length of the gut (23). There were no growth differences between *S.* Cerro isolates grown aerobically in either LB broth (Figure 1a) or M9-glucose minimal media (Figure 1b). *S*. Cerro isolates also grew with similar kinetics in LB broth in anaerobic conditions (Figure 1c). S. Typhimurium and *S.* Cerro grew similarly in all conditions tested *in vitro*, allowing us to compare *S*. Cerro isolates with the heavily studied organism, *S*. Typhimurium, in infection-relevant conditions *in vitro*.

**Figure 1:**
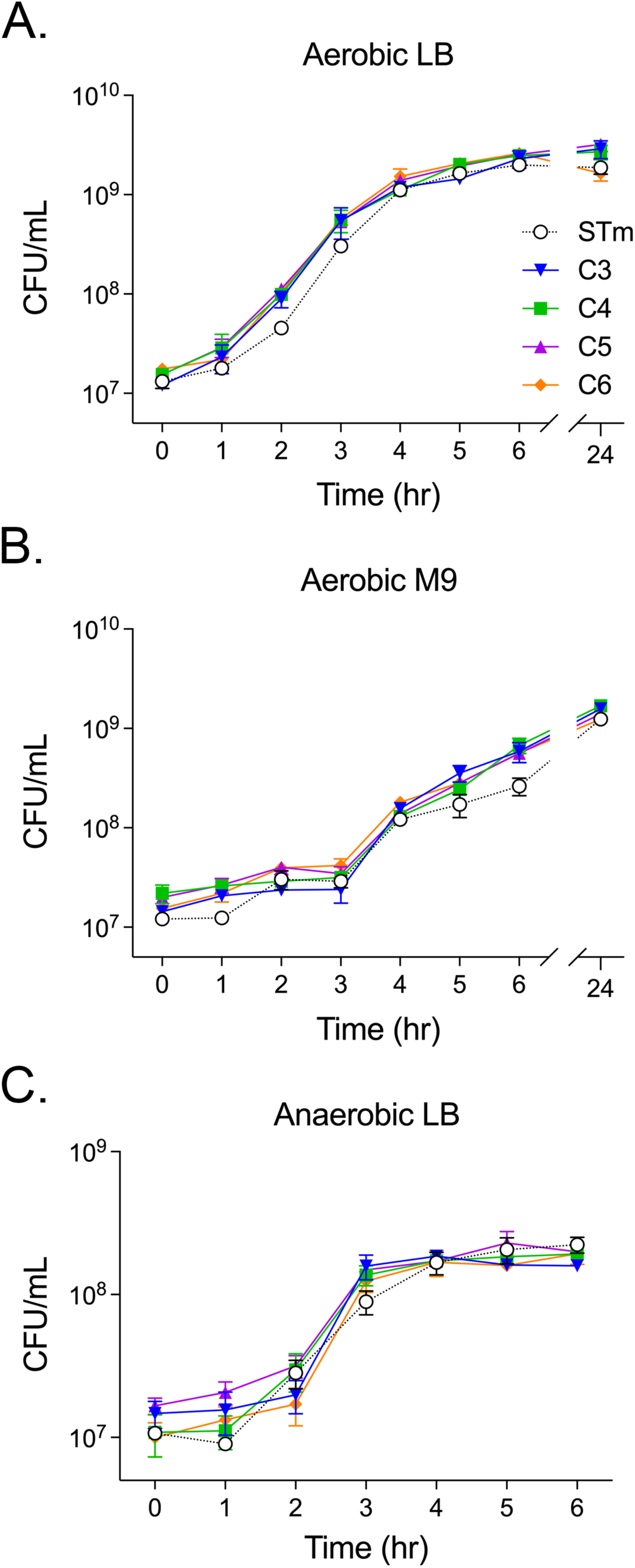
*S.* Cerro isolates grow similarly *in vitro.* Overnight cultures were diluted into LB (A) or M9 (B) broth and incubated aerobically at 37°C for 24 hours. (C) Pelleted overnight cultures were transferred into an anaerobic chamber, diluted into pre-reduced LB broth, and incubated anaerobically at 37°C for 6 hours. At the indicated times, an aliquot of bacteria was serially diluted and plated on LB agar to determine CFU/mL. Data represent the mean ± standard error of the mean (SEM) from 3 independent experiments. STm - *Salmonella* Typhimurium

Flagella-mediated motility is a virulence determinant for enteric infections with non-typhoidal *Salmonella* spp. (24). Therefore, we sought to establish the swimming motility of our *S*. Cerro isolates *in vitro*. Given a previous study indicated that *S.* Cerro has reduced motility (17), we hypothesized the *S.* Cerro isolates would have low to absent motility. We found that swimming motility is present among our *S.* Cerro isolates *in vitro*, although it appears to be less motile than *S*. Typhimurium *in vitro* (Figure 2a, b).

**Figure 2:**
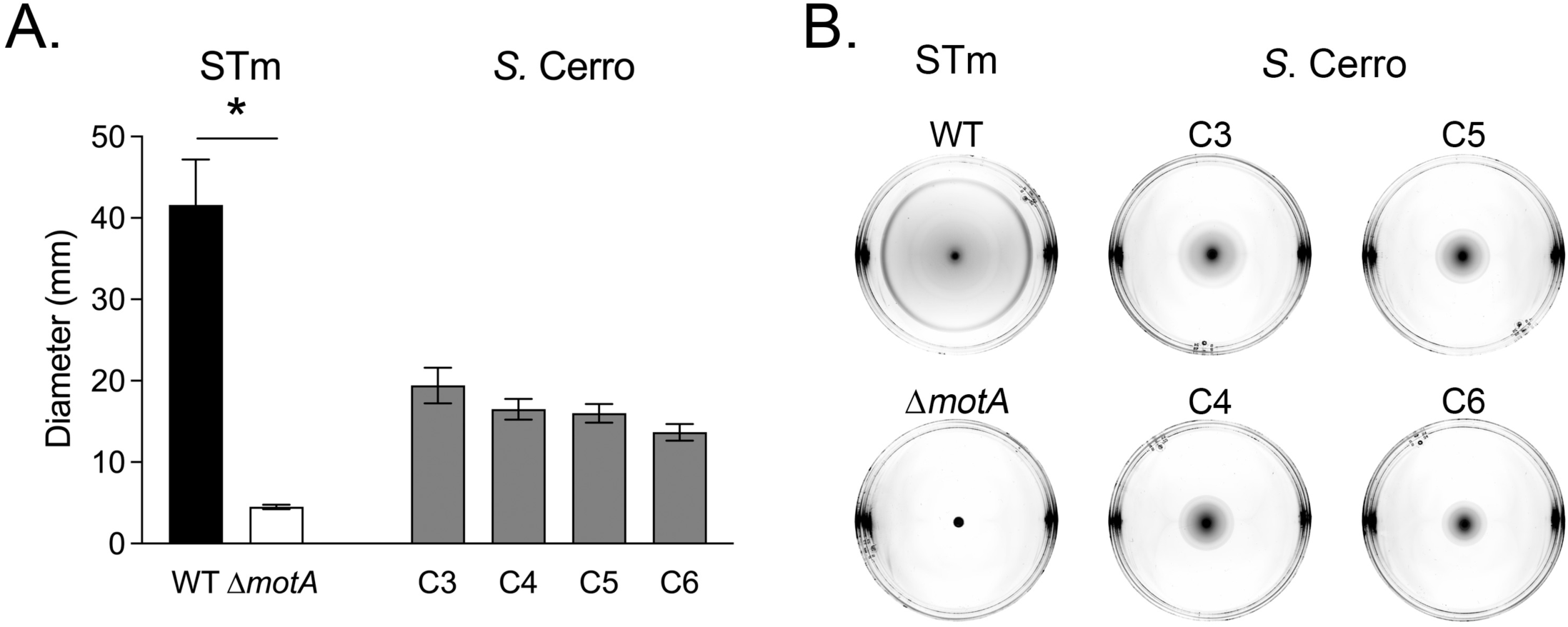
*S.* Cerro isolates have similar swimming motility *in vitro*. Normalized overnight cultures were spotted onto swimming agar plates in duplicate. Plates were incubated at 37°C for 6 hours. (A) Bars represent the mean ± SEM average diameter (mm) from 3 independent experiments. (B) Representative photographs of individual swimming plates are shown. Differences (*) between strains of the same serotype were determined by one-way ANOVA with Šídák’s correction for multiple comparisons (P<0.05)

Biofilm production contributes to environmental persistence of pathogens (25, 26). Thus, we determined the biofilm-forming capacity of our *S*. Cerro isolates in liquid culture. Three of four *S.* Cerro isolates produced robust biofilms (Figure 3a). Isolate C3 produced significantly less biofilm than the other three *S.* Cerro isolates, despite having similar planktonic cell density (Figure 3b). The planktonic cell density of isolate C6 was significantly less than isolate C5 in no-salt LB media, but this did not have measurable impact on biofilm production in liquid culture. These data demonstrate significant variation in biofilm formation in liquid culture for the temporally related *S.* Cerro isolates.

**Figure 3:**
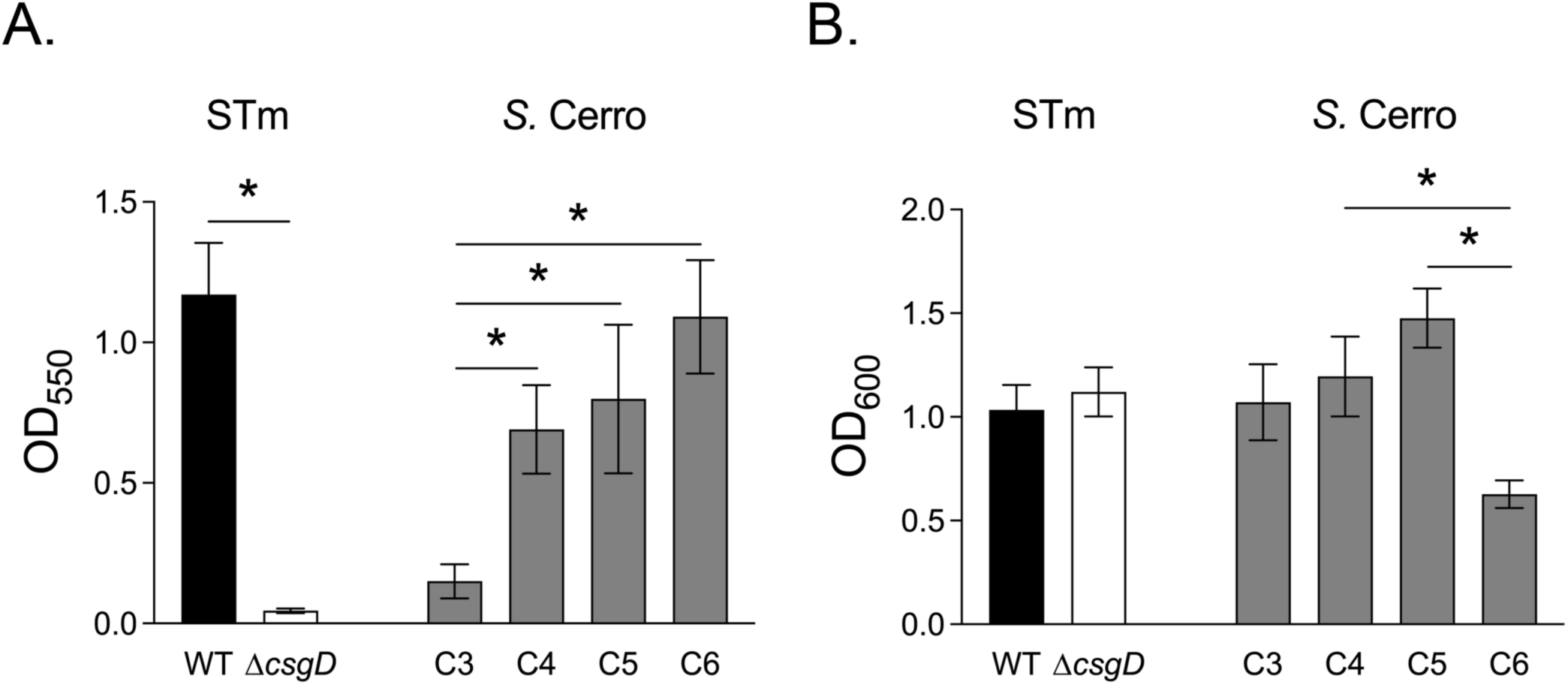
Biofilm formation varies between *S.* Cerro isolates. Overnight cultures were diluted into no-salt LB broth and were incubated at 25°C standing for 72 hours. Bars represent the mean ± SEM of (A) biofilm-associated cells as assessed by crystal violet staining or (B) planktonic cell density from 6 independent experiments. Differences (*) between strains of the same serotype were determined by one-way ANOVA with Šídák’s correction for multiple comparisons on log transformed data (P<0.05).

### Susceptibility of S. Cerro isolates to host-related stressors

Next, we sought to establish the sensitivity of *S.* Cerro isolates to stress conditions found within the mammalian gastrointestinal tract. We first tested acidic pH and bile salts as these are stressors encountered by foodborne pathogens in the abomasum (glandular stomach) and small intestine (27). All *S.* Cerro isolates were highly sensitive to acid stress (Figure 4a), with a mean percent survival +/- standard deviation of 0.52% +/- 0.27 for C3, 1.55% +/- 0.68 for C4, 3.23% +/- 1.19 for C5, and 2.73% +/- 1.93 for C6. The sensitivity to acid stress amongst *S*. Cerro isolates was similar to that of the acid stress-sensitive *S.* Typhimurium Δ*fur* mutant (6.57% +/- 5.29) (28), and survival was an order of magnitude lower than that of the WT *S.* Typhimurium (38.13% +/- 42.95).

**Figure 4:**
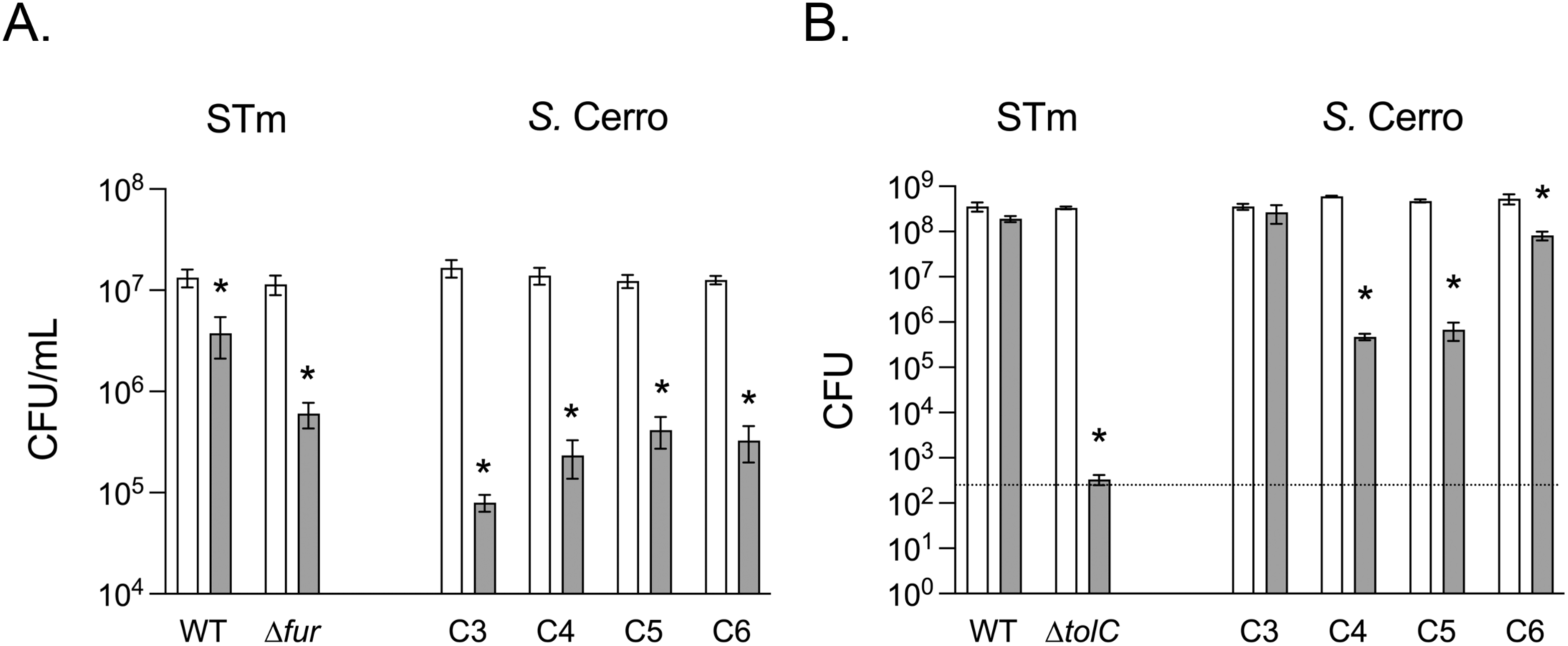
*S.* Cerro isolates are uniformly sensitive to acid and variably sensitive to bile salt exposure. (A) Overnight cultures were diluted into LB at pH 6.7 (white bars) or LB at pH 2.5 (gray bars). Viable bacteria were determined after 30 min exposure at 37°C by spot titer of serially diluted cultures. (B) Overnight cultures were diluted into LB and grown aerobically for 90 minutes at 37°C. Bacteria were normalized by OD_600_ and then serially diluted in PBS. Diluted cultures were spotted onto LB agar (white bars) or LB agar with 1% (w/v) deoxycholate (gray bars) and incubated at 37°C overnight. Viable bacteria were determined by spot titer. Bars represent mean ± SEM from 3 independent experiments and the dashed line represents the limit of detection. A two-way ANOVA with Šídák’s correction for multiple comparisons was used on log-transformed data to determine significant treatment effects (*) for each strain (p<0.05).

Unlike our observation for acid stress, the susceptibility of *S*. Cerro isolates to deoxycholate (DOC) was variable. Isolate C3 was resistant to deoxycholate, which was similar to what we observed for *S.* Typhimurium (Figure 4b). On the other hand, isolates C4, C5, and C6 lost significant viability when grown in the presence of DOC. The *S.* Typhimurium Δ*tolC* mutant was highly sensitive to DOC, consistent with prior reports (29). Together, these data suggest that gastric acid is likely to present a substantial barrier for *S.* Cerro intestinal colonization, but bile acids may not be effective antimicrobial agents against all *S.* Cerro isolates.

*Salmonella* can resist the toxic effects of bile salts by exporting them outside of the cell using the AcrAB-TolC efflux pump (29). Increased expression of the AcrAB-TolC efflux pump contributes to antimicrobial resistance (30). Because we demonstrated variable susceptibility of *S.* Cerro isolates to bile salt exposure, we hypothesized that the bile-sensitive *S.* Cerro isolates would be more sensitive to antibiotics than the bile-resistant isolates. We tested the sensitivity of *S.* Cerro isolates to five antibiotics commonly used for treatment of dairy calf diseases. We found that all *S.* Cerro isolates were susceptible to ampicillin, enrofloxacin, ceftiofur, and tetracycline and all were resistant to erythromycin *in vitro* (Table 2). Therefore, enhanced bile sensitivity in isolates C4, C5, and C6 did not impact sensitivity to antibiotics.

**Table 2.** Antibiotic susceptibility testing.

| Isolate | AMP* | Int. | ERYTH* | Int. | ENRO* | Int. | TET* | Int. | EXC* | Int. |
| --- | --- | --- | --- | --- | --- | --- | --- | --- | --- | --- |
| C3 | 28 (1) | S | 10.7 (1.2) | R | 34.7 (1.5) | S | 30 (0) | S | 30.3 (0.6) | S |
| C4 | 28.7 (1.5) | S | 11 (0) | R | 34.3 (0.6) | S | 32.7 (2.5) | S | 30.7 (1.2) | S |
| C5 | 28 (0) | S | 10.7 (0.6) | R | 33.7 (1.2) | S | 33 (1.7) | S | 31.3 (2.3) | S |
| C6 | 29.7 (1.2) | S | 10 (1.7) | R | 33.7 (1.5) | S | 20.3 (0.6) | S | 30.7 (1.2) | S |

\*Zone of inhibition (mm), mean (+/- SD)
AMP – ampicillin; ERYTH – erythromycin; ENRO – enrofloxacin; TET – tetracycline; EXC – ceftiofur; Int. – interpretation, S – susceptible, R – resistant

*Salmonella* are exposed to reactive oxygen and nitrogen species derived from the innate immune response to infection (31). Genomic studies suggest many *S.* Cerro isolates have lost the D-alanine amino acid transporter (11), which mediates resistance to reactive oxygen species (32). Therefore, we hypothesized that *S.* Cerro strains would be susceptible to the reactive oxygen species hydrogen peroxide and hypochlorite *in vitro*. We found that all *S.* Cerro isolates were highly susceptible to both hydrogen peroxide and hypochlorite (Figures 5a-b) at a dose which did not cause significant viability loss to *S*. Typhimurium WT. As anticipated, the *S.* Typhimurium Δ*recA* and Δ*yjiE* mutants were sensitive to hydrogen peroxide and hypochlorite, respectively (Figures 5a-b). Given the heightened sensitivity of *S*. Cerro isolates to reactive oxygen species, we hypothesized that *S*. Cerro isolates would also be sensitive to reactive nitrogen species *in vitro*. We observed no significant impact of reactive nitrogen species on the viability of *S*. Cerro isolates, similar to what we observed for *S.* Typhimurium (Figure 5c). Taken together, these data indicate *S.* Cerro isolates are poorly able to withstand some innate immune defenses encountered within the mammalian gastrointestinal tract.

**Figure 5:**
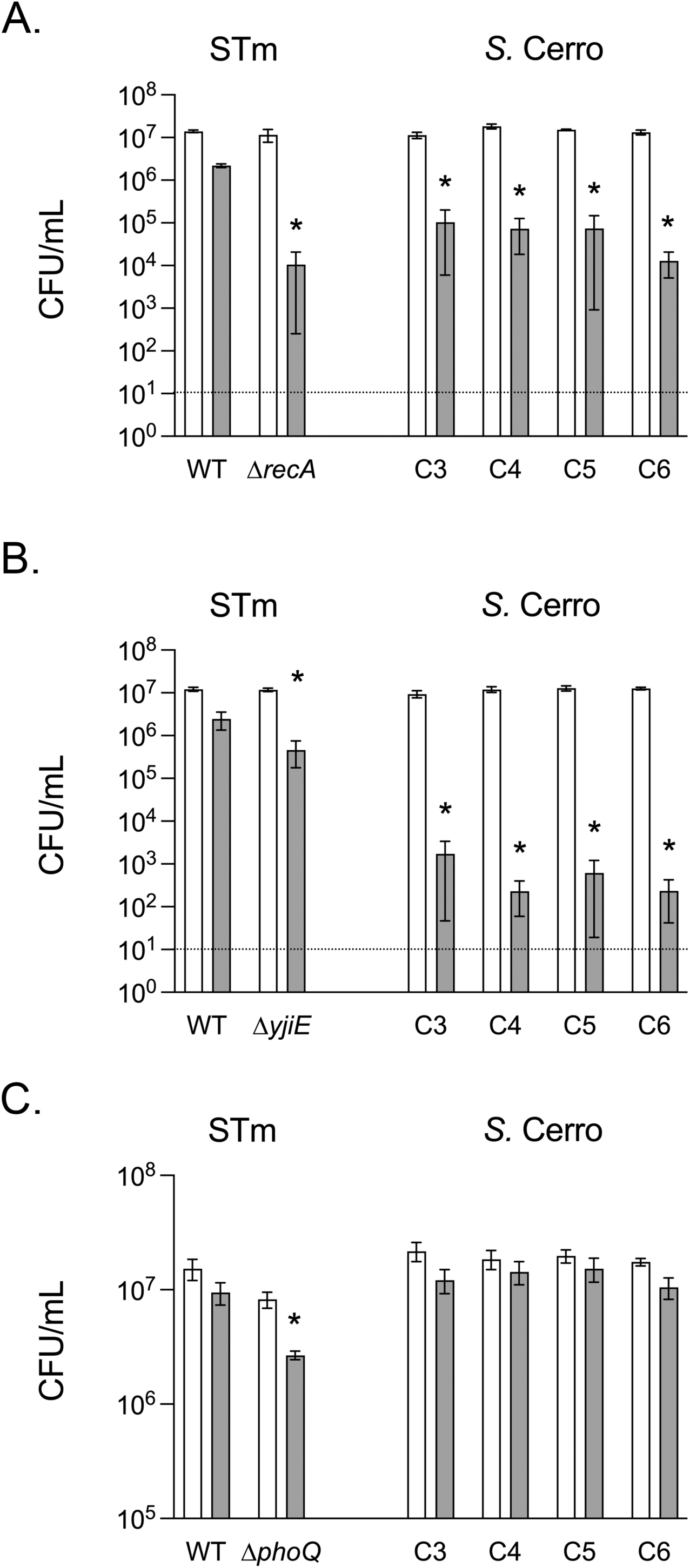
*S.* Cerro isolates are sensitive to reactive oxygen species and are resistant to reactive nitrogen species. Overnight cultures were diluted to ∼1x10^7^ CFU/mL in PBS. (A) Bacteria were treated with 10 mM hydrogen peroxide or not treated for 60 minutes, (B) sodium hypochlorite (5 μg/ml) for 30 minutes, or (C) spermine NONOate (250 μM) for 6 hr. Viable cells remaining after exposure were enumerated in serially diluted cultures by spot titer or plating. Gray bars indicate treated cultures, white bars are untreated control cultures and the dashed line represents the limit of detection. Bars indicate mean ± SEM from 3 (A), 5 (B) or 4 (C) independent experiments. A two-way ANOVA with Šídák’s correction for multiple comparisons was used on log-transformed data to determine significant treatment effects (*) for each strain (p<0.05).

### Fitness of S. Cerro isolates in the bovine intestine

*S.* Cerro isolates have disruptions to the T3SS1 effector *sopA* and reduced invasiveness *in vitro* (17), leading us to hypothesize that our *S*. Cerro isolates would be defective for colonization of bovine intestinal tissue. We performed competitive infections between two *S*. Cerro isolates and the virulent *S*. Typhimurium using the bovine ligated intestinal loop model. We tested isolate C3, as this isolate differs from the other *S.* Cerro isolates in biofilm formation and bile salt sensitivity, and isolate C5, which is representative of the other isolates *in vitro*. We harvested tissues 2-hours post-infection to measure tissue invasion before the onset of substantial neutrophilic inflammation (33) and found that both *S*. Cerro isolates were defective for intracellular colonization when compared with *S*. Typhimurium (Figure 6). In contrast, *S*. Typhimurium and *S*. Cerro isolates are equally fit in luminal fluid. These data support the hypothesis that *S.* Cerro isolates are attenuated for colonization of bovine intestinal tissue when compared with *S.* Typhimurium.

**Figure 6:**
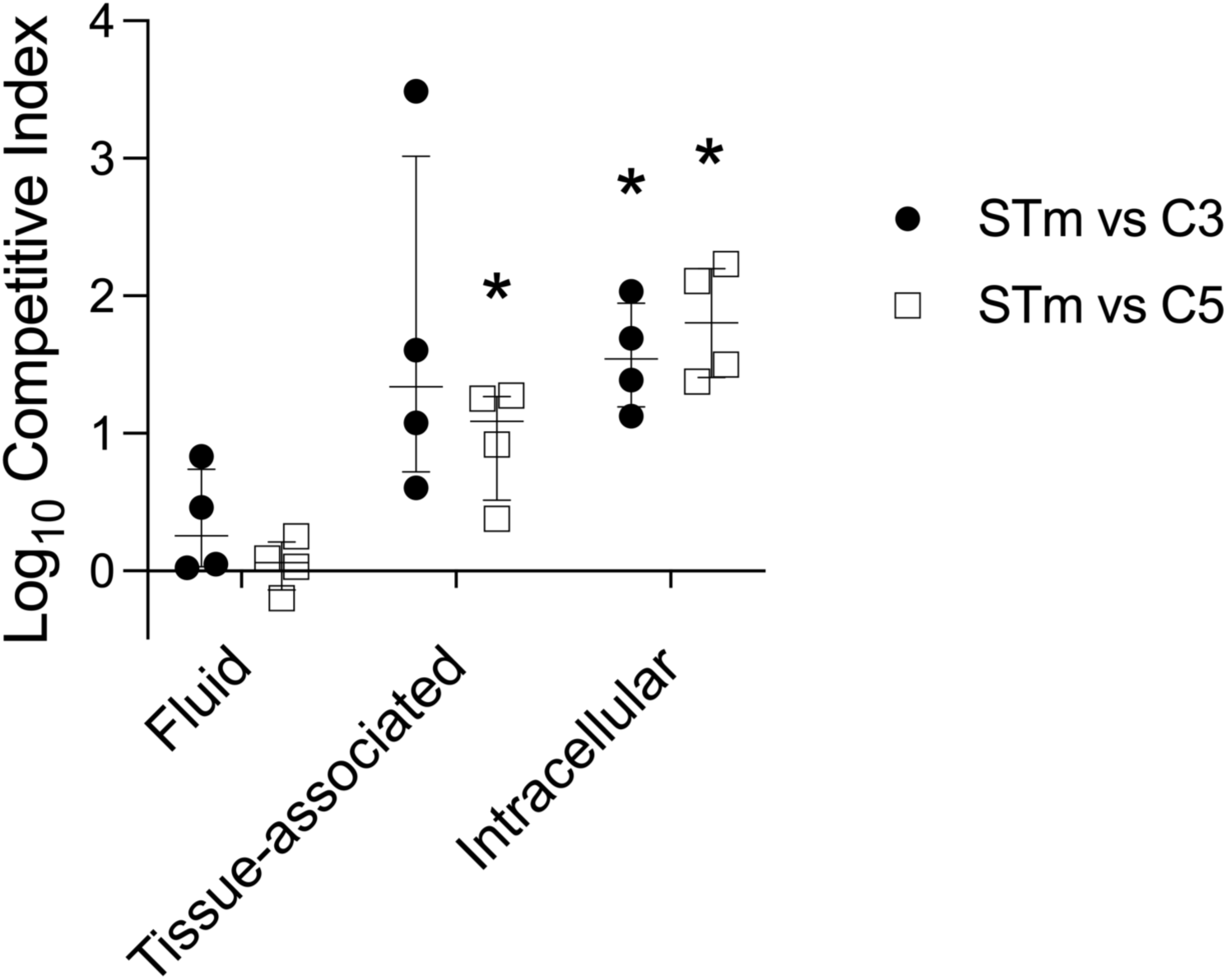
*S.* Cerro isolates poorly colonize bovine intestinal tissue. Ligated intestinal segments were inoculated with ∼1x10^9^ CFU of an equal mixture of *S.* Typhimurium strain ST4/74 (JE1909) and *S.* Cerro isolate C3 (JE2830) or C5 (JE2832). Calves were euthanized 2-hours post-infection and intestinal segments were harvested. Luminal fluid and intestinal tissue were separated. Intestinal tissue was split in half and was either untreated (tissue-associated) or treated with gentamicin (intracellular). Competitive index (CI) was the ratio of S. Typhimurium to S. Cerro isolate after infection divided by that ratio in the inoculum. Data points represent a single animal with median +/- interquartile range indicated. Asterisks (*) denote significant difference (p<0.05) in log_10_ competitive index as determined by unpaired t-test with Holm-Šídák’s correction for multiple comparisons.

## Discussion

In this study we evaluated infection-relevant *in vitro* phenotypic characteristics of four temporally-related *S.* Cerro strains isolated from neonatal calves on a dairy farm in Wisconsin. All of our *S.* Cerro isolates were highly sensitive to acid stress and reactive oxygen species but resistant to reactive nitrogen species *in vitro*. We observed phenotypic variation between isolates in biofilm formation and sensitivity to bile salts. Two *S*. Cerro isolates were defective for colonization of intestinal tissue when compared with *S.* Typhimurium during acute enteric infection of the bovine host. Our data suggest that *S.* Cerro is likely to be sensitive to numerous antibacterial killing mechanisms employed by the innate immune system during enteric salmonellosis.

*Salmonella* is commonly isolated from the feces of dairy cows and calves as well as from the farm environment (3). *S.* Cerro is an emerging *Salmonella* serotype isolated from cattle, but it is unclear whether it is a cause of concern for bovine health. Although *S*. Cerro can be isolated from feces of both apparently healthy and diseased cattle, the correlation of *S.* Cerro infection with disease varies between studies (8, 9). Here, we observed mild disease in neonatal calves naturally infected with *S*. Cerro within the first day of life. All four calves had intermittent, mild fever, with no other evidence of systemic disease. We were surprised to observe very few signs of systemic disease in *S*. Cerro-infected neonatal calves given that younger animals are more susceptible to invasive *Salmonella* infections than older animals (34–36). At post-mortem sampling, we documented *S.* Cerro in the mesenteric lymph nodes of two calves, suggesting the organism could survive within extra-intestinal tissues. Our *in vitro* data demonstrated S. Cerro isolates were resistant to reactive nitrogen species *in vitro,* suggesting that S. Cerro can withstand nitrosative stress encountered within tissue macrophages during prolonged infection (31, 37, 38). One limitation is that we are unable to establish whether the colonization of mesenteric lymph nodes resulted from a persistent or transient infection and whether the two animals without observed mesenteric lymph node colonization at post-mortem had cleared a prior infection. Our combined observations provide evidence that *S.* Cerro is capable of colonizing lymph nodes while eliciting only mild signs of disease in neonatal calves. Whether long-term lymph node colonization impacts growth or other health metrics in cattle remains unknown.

Diarrhea is a prominent feature of non-typhoidal *Salmonella* infections but is less common during infections with host-adapted serotypes. The T3SS1 and its effector proteins mediate invasion into intestinal epithelial cells, leading to a release of inflammatory cytokines and chemokines and subsequent neutrophil influx which causes inflammatory diarrhea. A common mutation observed in bovine *S.* Cerro isolates results in a premature stop codon in the T3SS1 effector protein SopA, which disrupts the ubiquitin ligase domain of the protein (10, 11). SopA contributes to fluid accumulation and neutrophil influx in bovine and *in vitro* infection models and a *S.* Typhimurium Δ*sopA* mutant is less virulent during oral infection in calves (14, 39). Disruption to the coding sequence of *sopA* is observed in the human host-restricted *Salmonella* serotype Typhi and is hypothesized to contribute to the reduction in gastrointestinal disease manifestation during Typhoid fever as compared with enteric salmonellosis (40). Furthermore, some *S.* Cerro isolates have reduced expression of the T3SS1 when compared with the broad host range serotype Typhimurium and the bovine host-adapted serotype Dublin (15, 16). Consistent with altered gastrointestinal colonization capacity of our *S*. Cerro isolates, only two of four calves with natural infection had mild to moderate severity diarrhea at any point during observation in our study. However, we did not test the calves for other enteric pathogens, so we are unable to definitively determine whether *S.* Cerro infection was the cause of diarrhea in these calves. The small number of animals with diarrhea, short duration, and low severity all are consistent with altered T3SS1 activity *in vivo*.

We found that *S.* Cerro isolates C3 and C5 had fitness defects when compared with *S.* Typhimurium in acute infection of the calf intestine. We observed reduced S. Cerro in association with intestinal tissue and in a gentamicin-protected intracellular niche when compared with S. Typhimurium. Both isolates were recovered from luminal contents in similar amounts, suggesting the two serotypes can co-exist in the gut lumen. A competitive disadvantage in intestinal tissue suggests *S*. Cerro isolates may be: i) impaired in invading intestinal tissue, or ii) killed by antimicrobial defenses at the intestinal epithelium. Decreased swimming motility among *S.* Cerro isolates, as compared to *S*. Typhimurium, could impact progression through the mucus layer to the intestinal epithelium, which is necessary to initiate invasion (24, 41). Alterations to T3SS1 expression or function could also impact intracellular S. Cerro numbers (42). We chose to compare infection dynamics between serotypes at an early time in infection, prior to the development of severe neutrophilic inflammation (33). By testing competitive fitness before inflammation, it is less likely that heightened sensitivity of *S*. Cerro isolates to inflammation-derived reactive oxygen species play a role in the reduced tissue colonization. Overall, the poor colonization of intestinal tissue by the *S.* Cerro isolates, in competition with *S.* Typhimurium, lends further support to the hypothesis that *S.* Cerro is attenuated for virulence in the bovine host.

We found that *S.* Cerro isolates were more sensitive to oxidative stress *in vitro* than *S.* Typhimurium. Lineages of *S.* Cerro have documented deletions in a D-alanine importer which mediates protection against neutrophil D-amino acid oxidase-induced oxidative damage (11, 18). While these mutations are predicted to impact survival in the face of neutrophilic inflammation, a D-alanine importer mutant (Δ*dalS*) of *S.* Typhimurium survives as well as the wild-type when challenged with either hydrogen peroxide or hypochlorous acid *in vitro* (32). Although we did not determine whether our *S.* Cerro strains had intact operons encoding the D-alanine importer, the increased *in vitro* susceptibility of our S. Cerro isolates to inorganic oxidative stress suggests that other oxidative stress responses may be defective in our *S.* Cerro isolates. It is intriguing that animals with diarrhea (calves 5 and 6) did not have *S*. Cerro in lymph nodes at postmortem examination, whereas those that did not develop diarrhea (calves 3 and 4) had *S*. Cerro in lymph nodes at postmortem examination. These observational data suggest that reactive oxygen species produced by neutrophilic inflammation, manifested clinically as diarrhea, could reduce the survival of *S.* Cerro within the gastrointestinal tract and thus impair systemic colonization. Therefore, calves that are sub-clinically shedding may be at a higher risk of harboring *S.* Cerro in systemic tissues, such as in mesenteric lymph nodes, which can impact food safety (43).

Our data demonstrate a substantial degree of phenotypic variability in biofilm formation and sensitivity to bile salts among *S.* Cerro isolates obtained from calves on the same dairy farm during a 30-day interval. Isolate C3 produced minimal biofilm and was resistant to bile salts *in vitro*. While it is not clear whether the reduced biofilm formation capacity relates to bile salt resistance, our data suggest that the mechanism of bile salt resistance is unlikely to be related to enhanced activity of multidrug efflux pumps, as all isolates had uniform antibiotic susceptibility with resistance to only erythromycin. The ability of *Salmonella* spp. strains to form biofilms has been proposed as a conserved strategy to increase persistence (44). The capacity for biofilm formation may have evolved in *S.* Cerro isolates C4-C6, as increasing biofilm production capacity has been shown to evolve over time in *S.* Typhimurium strains persisting on swine farms (45). We hypothesize that the observed strain variability could indicate on-farm microevolution within *S*. Cerro isolates to improve persistence. Future studies correlating genotype phylogeny with phenotype using a larger sample size are required to determine the impact of strain variability on *S.* Cerro’s capacity for long term survival within a herd.

Overall, our data are consistent with the hypothesis that *S.* Cerro is attenuated for virulence in cattle. *S.* Cerro isolates have some phenotypic attributes similar to *S.* Typhimurium that could favor persistence within the bovine host and promote on-farm persistence, such as resistance to reactive nitrogen species and biofilm forming capacity. However, *S.* Cerro isolates are highly susceptible to stressors present in the intestine including acidic pH and bile salts, which leaves an open question as to how *S.* Cerro survives transit through the abomasum (glandular stomach) and small intestine to colonize the bovine host. We hypothesize that *S.* Cerro has evolved to thrive within a specific niche in the bovine host without causing overt disease, as our data suggest that *S*. Cerro is less likely to withstand the reactive oxygen species produced during neutrophilic enterocolitis. Subclinical infections of *S.* Cerro, however, may increase the likelihood of systemic colonization and potentially threaten food safety if there is lymph node contamination of harvested adipose or muscle tissue. Future studies should determine whether early colonization with *S.* Cerro leads to chronic shedding and subsequent systemic colonization of adult cattle and determine whether stimulation of innate immunity would improve clearance of *S.* Cerro to reduce the persistence within herds. Our work has potential to inform interventions to control *S.* Cerro on a farm such as the use of oxidative and acidic treatments to eradicate *S.* Cerro from environmental reservoirs.

## Materials and Methods

### Bacterial strains and growth conditions

*Salmonella* spp. strains used in the phenotypic analyses are listed in Table 3. *S*. Cerro isolates were derived from day 1 fecal samples from all four calves. Previously described S. Typhimurium strain ATCC 14028 gene deletions (46) were moved into S. Typhimurium strain ST4/74 by P22 bacteriophage-mediated transduction (47). Transductants were twice purified and the mutations were confirmed using colony PCR with external primers to the gene of interest. Bacterial cultures were grown on Luria Bertani-Miller (LB) agar or in LB broth (BD Difco^TM^) at 37°C with agitation (225 rpm) unless otherwise specified. Media was supplemented as necessary with streptomycin (100 mg/L), kanamycin (50 mg/L), and nalidixic acid (50 mg/L).

**Table 3.** Bacterial strains used in this study.

| Strain | Genotype | Reference |
| --- | --- | --- |
| JE1909 | *Salmonella enterica* serotype Typhimurium str. ST4/74, Strep<sup>r</sup> | (58) |
| JE2355 | JE1909 Δ*motA::kan*, Strep<sup>r</sup>, Kan<sup>r</sup> | This study |
| JE1945 | JE1909 Δ*csgD::kan*, Strep<sup>r</sup>, Kan<sup>r</sup> | This study |
| JE2419 | JE1909 Δ*yjiE::kan*, Strep<sup>r</sup>, Kan<sup>r</sup> | This study |
| JE2498 | JE1909 Δ*phoQ::kan*, Strep<sup>r</sup>, Kan<sup>r</sup> | This study |
| JE2375 | JE1909 Δ*recA::kan*, Strep<sup>r</sup>, Kan<sup>r</sup> | This study |
| JE2507 | JE1909 Δ*fur::kan*, Strep<sup>r</sup>, Kan<sup>r</sup> | This study |
| JE2510 | JE1909 Δ*tolC::kan*, Strep<sup>r</sup>, Kan<sup>r</sup> | This study |
| SR64 | C3; *Salmonella enterica* serotype Cerro isolated from Calf 3 (day 1 feces) | This study |
| SR65 | C4; *Salmonella enterica* serotype Cerro isolated from Calf 4 (day 1 feces) | This study |
| SR89 | C5; *Salmonella enterica* serotype Cerro isolated from Calf 5 (day 1 feces) | This study |
| SR97 | C6; *Salmonella enterica* serotype Cerro isolated from Calf 6 (day 1 feces) | This study |
| JE2830 | SR64 with spontaneous Nal<sup>r</sup> | This study |
| JE2832 | SR89 with spontaneous Nal<sup>r</sup> | This study |

### Calves with natural S. Cerro infection

All animal procedures were performed under approved University of Wisconsin-Madison Institutional Animal Care and Use Committee protocols (Protocols V006401 and V006249-R01).

From April 14, 2021 to May 07, 2021, four Holstein-Angus crossbred calves (2 heifers and 2 bulls) were obtained from a University-owned dairy farm within 2 hours of birth. Calves were removed from the farm and housed in a room isolated from all other animals in individual pens which allowed no nose-to-nose contact. Calves were each administered 300 g of IgG (Colostrx, AgriLabs) via esophageal feeder within the first day of life. Calves were then fed non-medicated milk replacer twice daily at ∼20% body weight and offered non-medicated starter pellet with whole screened corn, grass hay, and water *ad libitum*. Serum total protein was measured at 24 hours of age to evaluate passive transfer of immunity. Rectal temperatures were monitored twice daily for the first 3 days of age, and then once per day. Calf health was monitored twice daily by scoring fecal consistency, appetite and attitude using modified scoring systems (14, 48). Attitude and appetite were scored from 0 to 4 (0 = calf is bright, alert, responsive and rises readily to drink, 1 = calf is quiet but responsive and requires encouragement to rise and drink, 2 = calf is lethargic and requires repeated encouragement to rise and drink, 3 = calf is depressed, recumbent and unable to rise on its own, 4 = calf is recumbent and obtunded). Fecal consistency was scored from 1 to 5 (1 = normal feces, 2 = soft feces with loss of distinct conformation, 3 = loose feces with reduced solid matter, 4 = aqueous feces with markedly reduced solid matter, +/- fibrin, 5 = aqueous feces with no solid matter, +/- fibrin, +/- blood). Calves were sedated with xylazine (VetOne) and were humanely euthanized with Fatal-Plus pentobarbital sodium (Vortech Pharmaceuticals) at the ages indicated (Table 1).

### Salmonella *spp*. selective fecal and tissue diagnostic testing

Approximately 5 g of feces was collected on day 1 of life and at the indicated ages (Table 1) for selective culture and enrichment for *Salmonella* spp. Fecal samples were streaked onto MacConkey and Xylose lysine tergitol 4 (XLT4) agar plates (BD Difco^TM^) using sterile cotton swabs and were incubated for 48 hours at 37°C. Approximately 1 g of feces or 5 g of each tissue sample were added to tetrathionate broth (BD Difco^TM^) and were incubated overnight at 37°C, standing. A sample from the tetrathionate broth was streaked onto MacConkey and XLT4 agar and incubated at 37°C for 48h and inoculated into Rappaport-Vassiliadis (RV) broth (BD Difco^TM^) and incubated at 42°C for 24 hours. A sample from the RV broth was then streaked onto MacConkey and XLT4 agar and incubated at 37°C for 48 hours. Colonies that showed H_2_S production or were lactose negative were re-streaked onto MacConkey and XLT4 agar to verify the phenotype. If colonies were both H_2_S producing and lactose negative, they were streaked onto blood agar plates (BD Difco^TM^) and incubated for 24 hours at 37°C. Colonies from blood agar plates were used to inoculate EnteroScreen 4^TM^ agar stab (Hardy Diagnostics), which was utilized according to manufacturer instructions. Colonies were verified as *Salmonella* spp. if they were urease negative, lysine-decarboxylase positive, lysine deamination negative, negative for lactose fermentation, and H_2_S positive.

A selection of positive *Salmonella* spp. colonies was confirmed by PCR using primers external to the coding region for the *Salmonella*-specific gene, *invA* (*invA*outerFWD: 5’- TGAGGGTTCGCTATTAACCG-3’; *invA*outerREV: 5’-TGGCAATGCAAATAAATCCA-3’). Colony PCR was performed on *Salmonella*-positive colonies on blood agar using Taq DNA polymerase (New England Biolabs®) with an annealing temperature of 55°C and an extension time of 2.5 min. for a total of 35 cycles. The expected size of approximately 2700 bp was confirmed by agarose gel electrophoresis in positive colonies. Positive colonies, confirmed to have the *invA* gene, isolated from fecal samples on day 1 from all calves and from mesenteric lymph node (MLN) tissue samples from calves 3 and 4 were submitted to the University of Wisconsin Veterinary Diagnostic Laboratory for serotyping using the Kauffman-White classification scheme. All *S*. Cerro isolates derived from fecal and tissue samples were preserved in 30% glycerol stocks at -80°C.

### Calf Ligated Intestinal Loop Surgery and Infection

For calf surgical infections, 4 male Holstein or Angus-Holstein cross-bred calves were obtained from a university-owned dairy farm within 2 days of birth. Calves were housed in AALAC-approved housing either individually or with nose-to-nose contact with one other calf. Calves were examined by a veterinarian to assess clinical condition upon arrival. Calves were either fed colostrum on-farm or a commercial colostrum replacer following transport (Colostrx, AgriLabs), depending on age at time of acquisition, with adequate passive transfer of immunity established 1-2 days after birth by serum total protein measurement. Calves were fed a commercial milk replacer at ∼20% body weight per day with non-medicated starter pellet, grass hay and water available *ad libitum*. Selective fecal cultures were performed at least twice weekly to establish *Salmonella* shedding status prior to surgery. *Salmonella* serogroups group B and K were not detected in any fecal cultures however all calves were fecal culture positive for *Salmonella* serogroup C confirmed by the Wisconsin Veterinary Diagnostic Laboratory. All calves had at least one negative fecal culture for *Salmonella* prior to surgery.

For surgical infections, bacteria were grown overnight at 37°C with agitation in LB broth. Overnight cultures were subcultured 1:100 into LB broth and incubated for approximately 4 hours at 37°C with agitation. Bacteria were washed twice in PBS and *S*. Typhimurium and *S*. Cerro isolates were mixed in a 1:1 ratio based on optical density (OD_600_).

Between 3 and 6 weeks of age, calves were anesthetized for ligated intestinal loop surgeries with minor modifications (20). The skin was desensitized with lidocaine hydrochloride (2%) before making the incision. Segments of the intestine were occluded in the aboral jejunum and ileum. The length of each intestinal segment was measured prior to inoculation with ∼10^9^ CFU containing an equal mixture of the indicated bacterial strains. Infected intestinal segments were harvested 2-hours post-infection and excised to collect luminal contents and tissue. Each tissue segment was washed twice in PBS to remove extracellular debris and unbound bacteria and then divided in half, with one segment treated with gentamicin (50 µg/mL) for 30 minutes at 37°C to quantify intracellular bacteria, while the remaining half was processed to quantify tissue-associated bacteria. After gentamicin treatment, tissues were washed twice with PBS. All samples were then placed in sterile PBS, homogenized, diluted, and plated to determine CFU/mL. Competitive index was calculated as the ratio of S. Typhimurium/S. Cerro after infection compared with the same ratio before infection.

### Growth curves

For evaluation of aerobic growth, overnight cultures were diluted 1:100 into LB or M9-glucose minimal media (48 mM Na_2_HPO_4_, 22 mM KH_2_PO_4_, 9 mM NaCl, 19 mM NH_4_Cl, 0.1 mM CaCl_2_, and 2 mM MgSO_4_ with 0.2% (wt/vol) glucose) and grown at 37°C with agitation. For anaerobic growth curves, overnight cultures were pelleted and supernatant decanted. Cell pellets were transferred into an anaerobic chamber (BactronEZ, Shel Lab) and resuspended in LB media that was pre-reduced for at least 18 hours. Bacteria were diluted 1:100 into pre-reduced LB media and incubated at 37°C. At the indicated times, bacteria were removed, serially diluted, and plated for evaluation of CFU per mL. Each assay was performed on three independent occasions.

### Swimming motility

Swimming motility was evaluated on semi-solid agar plates containing 0.3% Difco Bacto agar (LB Miller base 25 g/L), with the *S*. Typhimurium amotile mutant, *ΔmotA,* used as a negative control for the assay (49). Overnight cultures were normalized by optical density at 600 nm (OD_600_). Each strain was spotted (5 μL) onto the center of individual swimming agar plates in duplicate and plates were incubated at 37°C for 6 hours. The diameter of cell spread was recorded in 2 dimensions and averaged within a plate; data from two technical repeats were averaged for analysis. Photographs were taken of each plate (ChemiDoc MP, BioRad). The assay was performed on three independent occasions.

### Biofilm production

Overnight cultures were diluted 1:100 into no-salt LB broth in 5 mL polystyrene test tubes. The *S*. Typhimurium *ΔcsgD* mutant, which does not produce biofilm (50), was used as a negative control for the assay. Cultures were grown for 72 hours standing at 25°C. At 72 hours’ growth, the OD_600_ was measured on planktonic bacteria to monitor cell density. Biofilm was then quantified by crystal violet staining, as previously described (51). Briefly, washed tubes were stained with 0.1% crystal violet, which was solubilized in 30% acetic acid followed by measurement of OD_550_. Uninoculated media was used to determine the background for crystal violet staining. Each assay was performed on six independent occasions.

### Sensitivity to reactive oxygen and nitrogen species

Overnight cultures were diluted into phosphate buffered saline (PBS) to a concentration of approximately 10^7^ CFU/mL. To test sensitivity to reactive oxygen species (ROS), bacteria were treated with 5 μg/mL sodium hypochlorite (Sigma-Aldrich) for 30 minutes at 37°C or with 10 mM hydrogen peroxide (Sigma-Aldrich) for 60 minutes at 37°C (52). The *S*. Typhimurium mutants *ΔyjiE* and *ΔrecA* were included as positive controls for sensitivity to hypochlorite and hydrogen peroxide, respectively (53, 54). To test sensitivity to reactive nitrogen species (RNS), bacteria were treated with spermine-NONOate (250 μM) for 6 hours at 37°C (55). The *S*. Typhimurium Δ*phoQ* mutant was used as positive control for sensitivity to reactive nitrogen species (55). Viable cells remaining after each treatment were enumerated by spot titer or by plating diluted cultures. Sodium hypochlorite assays were performed on five independent occasions, spermine-NONOate assays were performed on four independent occasions and hydrogen peroxide assays were performed on three independent occasions.

### Sensitivity to acid shock

Overnight cultures were diluted 1:100 into LB pH 6.7 or into LB pH 2.5. The pH was adjusted in LB broth using 1N HCl. Viable cells remaining after 30 min of exposure were enumerated by spot titer or plating diluted cultures. The *S*. Typhimurium *Δfur* mutant was included as a control for acid sensitivity (28). Experiments were performed on three independent occasions.

### Sensitivity to deoxycholate

Overnight cultures were diluted 1:100 into LB broth and grown for 90 min at 37°C with agitation at 225 rpm. Bacteria were then normalized by OD_600_, serially diluted, and 4 μl was spotted onto LB agar or on LB agar with 1% (w/v) sodium deoxycholate (DOC, Thermo Fisher Scientific), as has been previously described (56). The plates were incubated at 37°C overnight and viable bacteria determined by spot titer. The *S*. Typhimurium *ΔtolC* mutant was used as a control for bile sensitivity (29). Experiments were performed on three independent occasions.

### Antibiotic susceptibility

Overnight cultures were normalized to an OD_600_ of 0.1 in sterile water. A lawn of each bacterial strain was spread onto large petri dishes containing Mueller-Hinton II agar (BD BBL^TM^) using sterile cotton swabs. Discs impregnated with erythromycin (BD; 15 μg), ampicillin (Microxpress; 10 μg), tetracycline (Microxpress; 30 μg), enrofloxacin (BD; 5 μg), and ceftiofur (BD; 30 μg) were placed on the bacterial lawns. An empty Whatman paper disc (GE Healthcare) was placed as a negative control in the center of the plate for each strain. Plates were incubated overnight at 37°C. The diameter of each zone of inhibition was measured at the widest point through the center of each disk. Interpretation of antibiotic sensitivity was based on updated Kirby-Bauer CLSI standards (57). Experiments were performed on three independent occasions.

### Statistical analysis and data availability

All raw data are available in supplemental table 1. The assay lower limit of detection minus one was used for graphical and statistical analyses for isolates with no viable cells detected when plated on 1% DOC or after ROS challenge. Shapiro-Wilk tests were used to establish whether data were normally distributed. CFU, CFU/mL, optical density, and competitive index data were log-transformed prior to analysis. For biofilm production and swimming motility, differences between strains within a given serotype were compared by one-way analysis of variance (ANOVA) with Šídák’s correction for multiple comparisons. For impacts of treatments on bacterial survival, a two-way ANOVA with Šídák’s correction for multiple comparisons was used to establish effects of a treatment within a given strain. Percent survival for acid stress was calculated as the CFU/mL upon acid treatment divided by the number of viable cells in the control condition, multiplied by 100. For competitive infections, competitive indices were log transformed and an unpaired t-test with Holm-Šídák’s correction for multiple comparisons used to determine the difference from zero. All analyses were performed using GraphPad Prism v10.1. Significance was set at p<0.05.

## Acknowledgements

This work was supported by start-up funds from the University of Wisconsin-Madison and Food Research Institute to JRE. JRE was supported, in part, by a US Department of Agriculture (USDA) National Institute of Food and Agriculture (NIFA) grant 2018-67017-27632. SMR was supported, in part, by a USDA NIFA Agriculture Food Research Initiative (AFRI) postdoctoral fellowship grant 2021-67012-35154. EC was supported by a Foundation for Food and Agricultural Research Veterinary Fellowship and the Science and Medicine Graduate Research Scholars Program at the University of Wisconsin-Madison. CLD was supported, in part, by a USDA NIFA AFRI predoctoral fellowship grant 2023-67011-40521.

## References

1. Kirk MD, Pires SM, Black RE, Caipo M, Crump JA, Devleesschauwer B, Döpfer D, Fazil A, Fischer-Walker CL, Hald T, Hall AJ, Keddy KH, Lake RJ, Lanata CF, Torgerson PR, Havelaar AH, Angulo FJ. 2015. World Health Organization Estimates of the Global and Regional Disease Burden of 22 Foodborne Bacterial, Protozoal, and Viral Diseases, 2010: A Data Synthesis. PLoS Med 12:e1001921.

2. Cummings KJ, Warnick LD, Elton M, Rodriguez-Rivera LD, Siler JD, Wright EM, Grohn YT, Wiedmann M. 2010. Salmonella enterica serotype Cerro among dairy cattle in New York: an emerging pathogen? Foodborne Pathog Dis 7:659–65.

3. Holschbach CL, Peek SF. 2018. Salmonella in Dairy Cattle. Vet Clin North Am Food Anim Pract 34:133–154.

4. Valenzuela JR, Sethi AK, Aulik NA, Poulsen KP. 2017. Antimicrobial resistance patterns of bovine Salmonella enterica isolates submitted to the Wisconsin Veterinary Diagnostic Laboratory: 2006-2015. J Dairy Sci 100:1319–1330.

5. Hong S, Rovira A, Davies P, Ahlstrom C, Muellner P, Rendahl A, Olsen K, Bender JB, Wells S, Perez A, Alvarez J. 2016. Serotypes and Antimicrobial Resistance in Salmonella enterica Recovered from Clinical Samples from Cattle and Swine in Minnesota, 2006 to 2015. PLoS One 11:e0168016.

6. Tewari D, Sandt CH, Miller DM, Jayarao BM, M’Ikanatha N M. 2012. Prevalence of Salmonella cerro in laboratory-based submissions of cattle and comparison with human infections in Pennsylvania, 2005-2010. Foodborne Pathog Dis 9:928–33.

7. Jones TF, Ingram LA, Cieslak PR, Vugia DJ, Tobin-D’Angelo M, Hurd S, Medus C, Cronquist A, Angulo FJ. 2008. Salmonellosis outcomes differ substantially by serotype. J Infect Dis 198:109–14.

8. Huston CL, Wittum TE, Love BC. 2002. Persistent fecal Salmonella shedding in five dairy herds. J Am Vet Med Assoc 220:650–5.

9. Chapagain PP, van Kessel JS, Karns JS, Wolfgang DR, Hovingh E, Nelen KA, Schukken YH, Grohn YT. 2008. A mathematical model of the dynamics of Salmonella Cerro infection in a US dairy herd. Epidemiol Infect 136:263–72.

10. Rodriguez-Rivera LD, Moreno Switt AI, Degoricija L, Fang R, Cummings CA, Furtado MR, Wiedmann M, den Bakker HC. 2014. Genomic characterization of Salmonella Cerro ST367, an emerging Salmonella subtype in cattle in the United States. BMC Genomics 15:427.

11. Kovac J, Cummings KJ, Rodriguez-Rivera LD, Carroll LM, Thachil A, Wiedmann M. 2017. Temporal Genomic Phylogeny Reconstruction Indicates a Geospatial Transmission Path of Salmonella Cerro in the United States and a Clade-Specific Loss of Hydrogen Sulfide Production. Front Microbiol 8:737.

12. Cohn AR, Orsi RH, Carroll LM, Liao J, Wiedmann M, Cheng RA. 2022. Salmonella enterica serovar Cerro displays a phylogenetic structure and genomic features consistent with virulence attenuation and adaptation to cattle. Frontiers in Microbiology 13.

13. Zhang S, Adams LG, Nunes J, Khare S, Tsolis RM, Baumler AJ. 2003. Secreted effector proteins of Salmonella enterica serotype typhimurium elicit host-specific chemokine profiles in animal models of typhoid fever and enterocolitis. Infect Immun 71:4795–803.

14. Zhang S, Santos RL, Tsolis RM, Stender S, Hardt W-D, Bäumler AJ, Adams LG. 2002. The Salmonella enterica Serotype Typhimurium Effector Proteins SipA, SopA, SopB, SopD, and SopE2 Act in Concert To Induce Diarrhea in Calves. Infect Immun 70:3843–3855.

15. Cohn AR, Orsi RH, Carroll LM, Chen R, Wiedmann M, Cheng RA. 2021. Characterization of Basal Transcriptomes Identifies Potential Metabolic and Virulence-Associated Adaptations Among Diverse Nontyphoidal Salmonella enterica Serovars. Front Microbiol 12:730411.

16. Salaheen S, Kim SW, Haley BJ, Van Kessel JAS. 2022. Differences between the global transcriptomes of Salmonella enterica serovars Dublin and Cerro infecting bovine epithelial cells. BMC Genomics 23:498.

17. Salaheen S, Sonnier J, Kim SW, Haley BJ, Van Kessel JAS. 2020. Interaction of Salmonella enterica with Bovine Epithelial Cells Demonstrates Serovar-Specific Association and Invasion Patterns. Foodborne Pathog Dis 17:608–610.

18. Osborne SE, Tuinema BR, Mok MC, Lau PS, Bui NK, Tomljenovic-Berube AM, Vollmer W, Zhang K, Junop M, Coombes BK. 2012. Characterization of DalS, an ATP-binding cassette transporter for D-alanine, and its role in pathogenesis in Salmonella enterica. J Biol Chem 287:15242–50.

19. Haneda T, Ishii Y, Danbara H, Okada N. 2009. Genome-wide identification of novel genomic islands that contribute to Salmonella virulence in mouse systemic infection. FEMS Microbiol Lett 297:241–9.

20. Elfenbein JR, Endicott-Yazdani T, Porwollik S, Bogomolnaya LM, Cheng P, Guo J, Zheng Y, Yang HJ, Talamantes M, Shields C, Maple A, Ragoza Y, Deatley K, Tatsch T, Cui P, Andrews KD, McClelland M, Lawhon SD, Andrews-Polymenis H. 2013. Novel Determinants of Intestinal Colonization of Salmonella enterica Serotype Typhimurium Identified in Bovine Enteric Infection. Infect Immun 81:4311–20.

21. Chaudhuri RR, Morgan E, Peters SE, Pleasance SJ, Hudson DL, Davies HM, Wang J, van Diemen PM, Buckley AM, Bowen AJ, Pullinger GD, Turner DJ, Langridge GC, Turner AK, Parkhill J, Charles IG, Maskell DJ, Stevens MP. 2013. Comprehensive Assignment of Roles for *Salmonella* Typhimurium Genes in Intestinal Colonization of Food-Producing Animals. PLoS Genet 9:e1003456.

22. Godden SM, Lombard JE, Woolums AR. 2019. Colostrum Management for Dairy Calves. Vet Clin North Am Food Anim Pract 35:535–556.

23. Friedman ES, Bittinger K, Esipova TV, Hou L, Chau L, Jiang J, Mesaros C, Lund PJ, Liang X, FitzGerald GA, Goulian M, Lee D, Garcia BA, Blair IA, Vinogradov SA, Wu GD. 2018. Microbes vs. chemistry in the origin of the anaerobic gut lumen. Proc Natl Acad Sci U S A 115:4170–4175.

24. Stecher B, Hapfelmeier S, Muller C, Kremer M, Stallmach T, Hardt WD. 2004. Flagella and chemotaxis are required for efficient induction of Salmonella enterica serovar Typhimurium colitis in streptomycin-pretreated mice. Infect Immun 72:4138–50.

25. Steenackers H, Hermans K, Vanderleyden J, De Keersmaecker SCJ. 2012. Salmonella biofilms: An overview on occurrence, structure, regulation and eradication. Food Research International 45:502–531.

26. Abdallah M, Benoliel C, Drider D, Dhulster P, Chihib NE. 2014. Biofilm formation and persistence on abiotic surfaces in the context of food and medical environments. Arch Microbiol 196:453–72.

27. Alvarez-Ordonez A, Begley M, Prieto M, Messens W, Lopez M, Bernardo A, Hill C. 2011. Salmonella spp. survival strategies within the host gastrointestinal tract. Microbiology 157:3268–81.

28. Hall HK, Foster JW. 1996. The role of fur in the acid tolerance response of Salmonella typhimurium is physiologically and genetically separable from its role in iron acquisition. J Bacteriol 178:5683–91.

29. Baucheron S, Mouline C, Praud K, Chaslus-Dancla E, Cloeckaert A. 2005. TolC but not AcrB is essential for multidrug-resistant Salmonella enterica serotype Typhimurium colonization of chicks. Journal of Antimicrobial Chemotherapy 55:707–712.

30. Prouty AM, Brodsky IE, Falkow S, Gunn JS. 2004. Bile-salt-mediated induction of antimicrobial and bile resistance in Salmonella typhimurium. Microbiology (Reading) 150:775–783.

31. Mastroeni P, Vazquez-Torres A, Fang FC, Xu Y, Khan S, Hormaeche CE, Dougan G. 2000. Antimicrobial Actions of the Nadph Phagocyte Oxidase and Inducible Nitric Oxide Synthase in Experimental Salmonellosis. II. Effects on Microbial Proliferation and Host Survival in Vivo. Journal of Experimental Medicine 192:237–248.

32. Tuinema BR, Reid-Yu SA, Coombes BK. 2014. Salmonella evades D-amino acid oxidase to promote infection in neutrophils. MBio 5:e01886.

33. Nunes JS, Lawhon SD, Rossetti CA, Khare S, Figueiredo JF, Gull T, Burghardt RC, Baumler AJ, Tsolis RM, Andrews-Polymenis HL, Adams LG. 2010. Morphologic and cytokine profile characterization of Salmonella enterica serovar typhimurium infection in calves with bovine leukocyte adhesion deficiency. Vet Pathol 47:322–33.

34. Nielsen LR, van den Borne B, van Schaik G. 2007. Salmonella Dublin infection in young dairy calves: transmission parameters estimated from field data and an SIR-model. Prev Vet Med 79:46–58.

35. Smith BP, Habasha F, Reina-Guerra M, Hardy AJ. 1979. Bovine salmonellosis: experimental production and characterization of the disease in calves, using oral challenge with Salmonella typhimurium. Am J Vet Res 40:1510–3.

36. Zhang K, Dupont A, Torow N, Gohde F, Leschner S, Lienenklaus S, Weiss S, Brinkmann MM, Ķhnel M, Hensel M, Fulde M, Hornef MW. 2014. Age-Dependent Enterocyte Invasion and Microcolony Formation by *Salmonella*. PLoS Pathog 10:e1004385.

37. Vazquez-Torres A, Jones-Carson J, Mastroeni P, Ischiropoulos H, Fang FC. 2000. Antimicrobial actions of the NADPH phagocyte oxidase and inducible nitric oxide synthase in experimental salmonellosis. I. Effects on microbial killing by activated peritoneal macrophages in vitro. J Exp Med 192:227–36.

38. Burton NA, Schurmann N, Casse O, Steeb AK, Claudi B, Zankl J, Schmidt A, Bumann D. 2014. Disparate impact of oxidative host defenses determines the fate of Salmonella during systemic infection in mice. Cell Host Microbe 15:72–83.

39. Wood MW, Jones MA, Watson PR, Siber AM, McCormick BA, Hedges S, Rosqvist R, Wallis TS, Galyov EE. 2000. The secreted effector protein of Salmonella dublin, SopA, is translocated into eukaryotic cells and influences the induction of enteritis. Cell Microbiol 2:293–303.

40. Valenzuela LM, Hidalgo AA, Rodriguez L, Urrutia IM, Ortega AP, Villagra NA, Paredes-Sabja D, Calderon IL, Gil F, Saavedra CP, Mora GC, Fuentes JA. 2015. Pseudogenization of sopA and sopE2 is functionally linked and contributes to virulence of Salmonella enterica serovar Typhi. Infect Genet Evol 33:131–42.

41. Horstmann JA, Zschieschang E, Truschel T, de Diego J, Lunelli M, Rohde M, May T, Strowig T, Stradal T, Kolbe M, Erhardt M. 2017. Flagellin phase-dependent swimming on epithelial cell surfaces contributes to productive Salmonella gut colonisation. Cell Microbiol 19.

42. Ibarra JA, Knodler LA, Sturdevant DE, Virtaneva K, Carmody AB, Fischer ER, Porcella SF, Steele-Mortimer O. 2010. Induction of Salmonella pathogenicity island 1 under different growth conditions can affect Salmonella–host cell interactions in vitro. Microbiology 156:1120–1133.

43. Arthur TM, Brichta-Harhay DM, Bosilevac JM, Guerini MN, Kalchayanand N, Wells JE, Shackelford SD, Wheeler TL, Koohmaraie M. 2008. Prevalence and characterization of Salmonella in bovine lymph nodes potentially destined for use in ground beef. J Food Prot 71:1685–8.

44. MacKenzie KD, Wang Y, Musicha P, Hansen EG, Palmer MB, Herman DJ, Feasey NA, White AP. 2019. Parallel evolution leading to impaired biofilm formation in invasive Salmonella strains. PLoS Genet 15:e1008233.

45. Tassinari E, Duffy G, Bawn M, Burgess CM, McCabe EM, Lawlor PG, Gardiner G, Kingsley RA. 2019. Microevolution of antimicrobial resistance and biofilm formation of Salmonella Typhimurium during persistence on pig farms. Sci Rep 9:8832.

46. Porwollik S, Santiviago CA, Cheng P, Long F, Desai P, Fredlund J, Srikumar S, Silva CA, Chu W, Chen X, Canals R, Reynolds MM, Bogomolnaya L, Shields C, Cui P, Guo J, Zheng Y, Endicott-Yazdani T, Yang H-J, Maple A, Ragoza Y, Blondel CJ, Valenzuela C, Andrews-Polymenis H, McClelland M. 2014. Defined Single-Gene and Multi-Gene Deletion Mutant Collections in *Salmonella enterica* sv Typhimurium. PLoS ONE 9:e99820.

47. Sternberg NL, Maurer R. 1991. Bacteriophage-mediated generalized transduction in Escherichia coli and Salmonella typhimurium, p 18–43. In Jeffrey HM (ed), Methods Enzymol, vol Volume 204. Academic Press.

48. McGuirk SM, Peek SF. 2014. Timely diagnosis of dairy calf respiratory disease using a standardized scoring system. Anim Health Res Rev 15:145–7.

49. Bogomolnaya LM, Aldrich L, Ragoza Y, Talamantes M, Andrews KD, McClelland M, Andrews-Polymenis HL. 2014. Identification of novel factors involved in modulating motility of Salmonella enterica serotype typhimurium. PLoS One 9:e111513.

50. Römling U, Rohde M, Olsén A, Normark S, Reinköster J. 2000. AgfD, the checkpoint of multicellular and aggregative behaviour in Salmonella typhimurium regulates at least two independent pathways. Molecular Microbiology 36:10–23.

51. Griewisch KF, Pierce JG, Elfenbein JR. 2020. Genetic determinants of Salmonella resistance to the biofilm inhibitory effects of a synthetic 4-oxazolidinone small molecule. Appl Environ Microbiol doi:10.1128/AEM.01120-20.

52. Hahn MM, Gunn JS. 2020. Salmonella Extracellular Polymeric Substances Modulate Innate Phagocyte Activity and Enhance Tolerance of Biofilm-Associated Bacteria to Oxidative Stress. Microorganisms 8.

53. Jo I, Kim D, No T, Hong S, Ahn J, Ryu S, Ha N-C. 2019. Structural basis for HOCl recognition and regulation mechanisms of HypT, a hypochlorite-specific transcriptional regulator. Proceedings of the National Academy of Sciences 116:3740–3745.

54. Karash S, Liyanage R, Qassab A, Lay JO, Jr., Kwon YM. 2017. A Comprehensive Assessment of the Genetic Determinants in Salmonella Typhimurium for Resistance to Hydrogen Peroxide Using Proteogenomics. Sci Rep 7:17073.

55. Bourret TJ, Liu L, Shaw JA, Husain M, Vazquez-Torres A. 2017. Magnesium homeostasis protects Salmonella against nitrooxidative stress. Sci Rep 7:15083.

56. Reynolds MM, Bogomolnaya L, Guo J, Aldrich L, Bokhari D, Santiviago CA, McClelland M, Andrews-Polymenis H. 2011. Abrogation of the twin arginine transport system in Salmonella enterica serovar Typhimurium leads to colonization defects during infection. PLoS ONE 6:e15800.

57. Humphries R, Bobenchik AM, Hindler JA, Schuetz AN. 2021. Overview of Changes to the Clinical and Laboratory Standards Institute Performance Standards for Antimicrobial Susceptibility Testing, M100, 31st Edition. J Clin Microbiol 59:e0021321.

58. Rankin JD, Taylor RJ. 1966. The estimation of doses of Salmonella typhimurium suitable for the experimental production of disease in calves. Vet Rec 78:706–7.

